# Deciphering the combinatorial interaction landscape

**DOI:** 10.1101/790543

**Authors:** Antonio Cappuccio, Shane T. Jensen, Boris Hartmann, Stuart C. Sealfon, Vassili Soumelis, Elena Zaslavsky

## Abstract

From cellular activation to drug combinations, the control of biological systems involves multiple stimuli that can elicit complex nonlinear interactions. To elucidate the functions and logic of stimulus interactions, we developed SAIL (Synergistic/Antagonistic Interaction Learner). SAIL uses a machine learning classifier trained to categorize interactions across a complete taxonomy of possible combinatorial effects. The strategy resolves the most informative interactions, and helps infer their functions and regulatory mechanisms. SAIL-predicted interaction mechanisms controlling key immune functions were experimentally validated. SAIL can integrate results from multiple datasets to derive general properties of how cells respond to multiple stimuli. Using public immunological datasets, we assembled a fine-grained landscape of ∼30000 interactions. Analysis of the landscape shows the context-dependent functions of individual modulators, and reveals a probabilistic algebra that links the separate and combined stimulus effects. SAIL is available through a user friendly interface to resolve the effect of stimulus and drug combinations.

## Introduction

Biological responses are shaped by the effects of multiple stimuli in complex environments, and treatment of many diseases involves combination therapies ^1,2^. A complex consequence of combination stimuli is the potential occurrence of nonlinear interactions that can dramatically alter the effects of individual treatments. Studies of the inflammatory microenvironment, for example, have identified various emergent effects of combination exposures ^3–5^. Although understanding such combinatorial effects is key to elucidating biological processes, new computational tools are still needed to dissect and interpret interactions between biological signals in *-omics* studies.

The typical combinatorial treatment experiment consists of *-omics* data generated in four conditions: 0 (control), stimulus X, stimulus Y, and the combination X+Y. A common way to understand interaction effects between X and Y is to identify non-additive responses induced by the combination X+Y, and to classify them as either synergistic -larger than additive- or antagonistic -smaller than additive-. Despite widespread use, this approach has limited resolution because it confounds interactions that are qualitatively different ^6^. For example, the conventional approach does not discriminate less common patterns of interaction with high biological significance, such as the emergence of a new response, from more frequent nominal interaction responses, such as non-additivity due to saturation effects. Furthermore, due to the lack of a satisfactory analysis framework and accessible tools, most studies are restricted to a specific combination of interest, providing only partial, fragmented insight. The narrow scope of current approaches may obscure general properties and principles that underlie the occurrence of combinatorial interactions.

Here, we present a comprehensive framework to map and interpret interaction effects within and across *-omics* combination treatments. The framework is based on a machine learning classifier trained to categorize gene responses in *-omics* combination treatments across a predefined, complete taxonomy of theoretically possible response patterns. Mapping the experimental gene responses into the appropriate element of the taxonomy resolves the most informative combinatorial effects, and facilitates the inference of coherent biological programs and of the underlying regulatory mechanisms. SAIL guided the identification of new cytokine interaction mechanisms in human dendritic cells, which we experimentally validated with neutralizing experiments.

Another major advance of SAIL is the capacity to integrate results from multiple *-omics* combination treatments and derive a broader understanding of how cells respond to combinations of stimuli. Using a compendium of public datasets, we assembled a fine-grained landscape comprising ∼30000 interactions from a variety of immune cells. Analysis of the landscape sheds new light on the context-dependent functions of individual modulators, and reveals a probabilistic algebra, a set of probabilistic rules underlying the integration process that link the separate and combined stimulus effects.

SAIL is available through a user friendly interface to resolve the combinatorial control of biological processes in public or user-generated dataset, and to assist the development of rational combination therapeutics.

## Results

### Overview of the SAIL framework

SAIL is a machine learning framework to map and interpret interaction effects from *-omics* combination treatment experiments comprising a vehicle control (denoted by 0), two individual stimuli (X, Y), and their combination (X+Y) (**Fig. 1a, left sub-panel**). Samples are harvested at specific timepoints after stimulation, and an -*omics* dataset, such as gene expression microarray or RNA-seq, is generated. The dataset is analyzed by a machine learning classifier previously trained to map gene responses across the predefined taxonomy of 123 possible response profiles (**Fig. S1**). These represent qualitatively different scenarios for the expression of a gene in the conditions 0, X, Y, X+Y ^6^. As we demonstrate, mapping the experimental gene responses into the appropriate element of the taxonomy resolves the most informative combinatorial effects, and facilitates the inference of coherent biological programs (**Fig. 1a, right sub-panel**).

**Figure 1.**
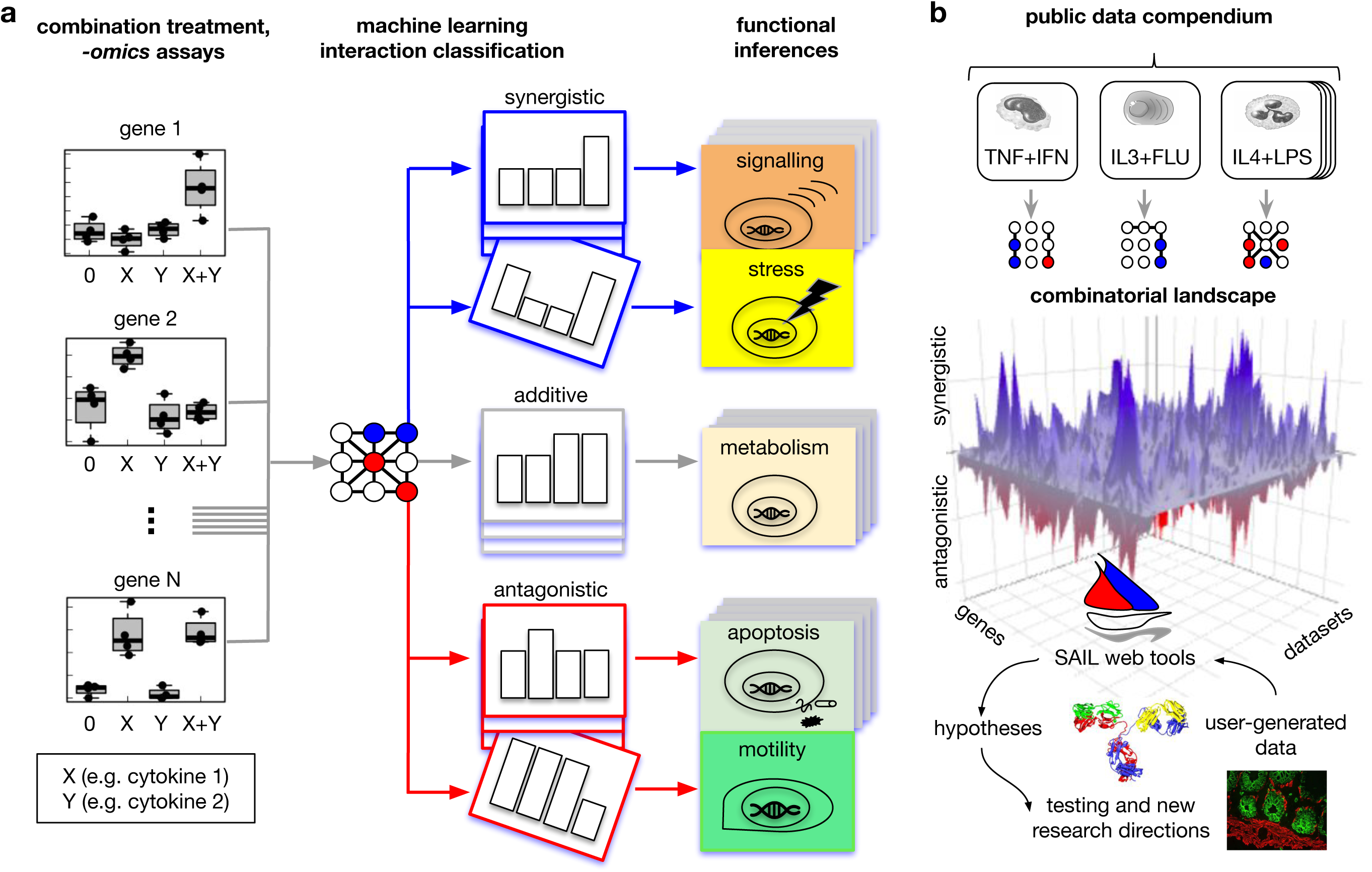
Overview of the SAIL framework. (**a**) SAIL (immune Synergistic/Antagonistic Interaction Learner) is a machine learning framework to decipher the effect of combination treatments. It takes as input -*omics* data from the four prototypical conditions of combination treatments: 0 (control), stimulus X, stimulus Y, and the combination X+Y (**left sub-panel**). The dataset is analyzed by a classifier trained to map each gene into a complete taxonomy of theoretically possible response profiles (**middle sub-panel**, see also **Fig. S1**). The taxonomy helps infer the functional role of different types of synergistic and antagonistic effects (**right sub-panel**). (**b**) A major advance of SAIL is the capacity to integrate results from multiple combination treatments. Using a compendium of publicly available immunological datasets (**right sub-panel**), we built a combinatorial landscape comprising ∼30000 interactions. Global analysis of the landscape and of user-generated data drive new hypotheses on the logic and functions of combinatorial interactions.

The use of an abstract taxonomy makes it possible to map interactions from multiple datasets onto a common reference space. Using public immunological datasets, we assembled ∼30,000 combinatorial interactions in an overall, fine-grained combinatorial landscape of immunity (**Fig. 1b**). The landscape is an information-rich object that reveals new aspects of the logic and functions of interactions, guiding new hypotheses.

### Machine learning-driven classification of combinatorial interactions

A key problem we address is how to classify noisy *-omics* data from combination treatments across the taxonomy of theoretical profiles. To solve this problem, we trained a machine learning classifier on an extensive set of simulated interaction profiles (**Fig. 2a**). Each interaction profile was simulated in multiple instances with variable group means for the conditions 0, X, Y, X+Y within a range of values consistent with the experimental data (**see Methods**). The statistical variability around the group means was assumed to be normally distributed. This assumption is widely held in the analysis of microarray data, and still applicable to RNA-seq data upon a suitable transformation ^7^. Given the uncertain noise level in the experimental data, we simulated three noise regimes: low, medium, and high (**Fig. 2a, see Methods**). The different noise levels were simulated by decreasing the effect size, defined in terms of the standardized means differences between the four conditions (**see Methods**).

**Figure 2.**
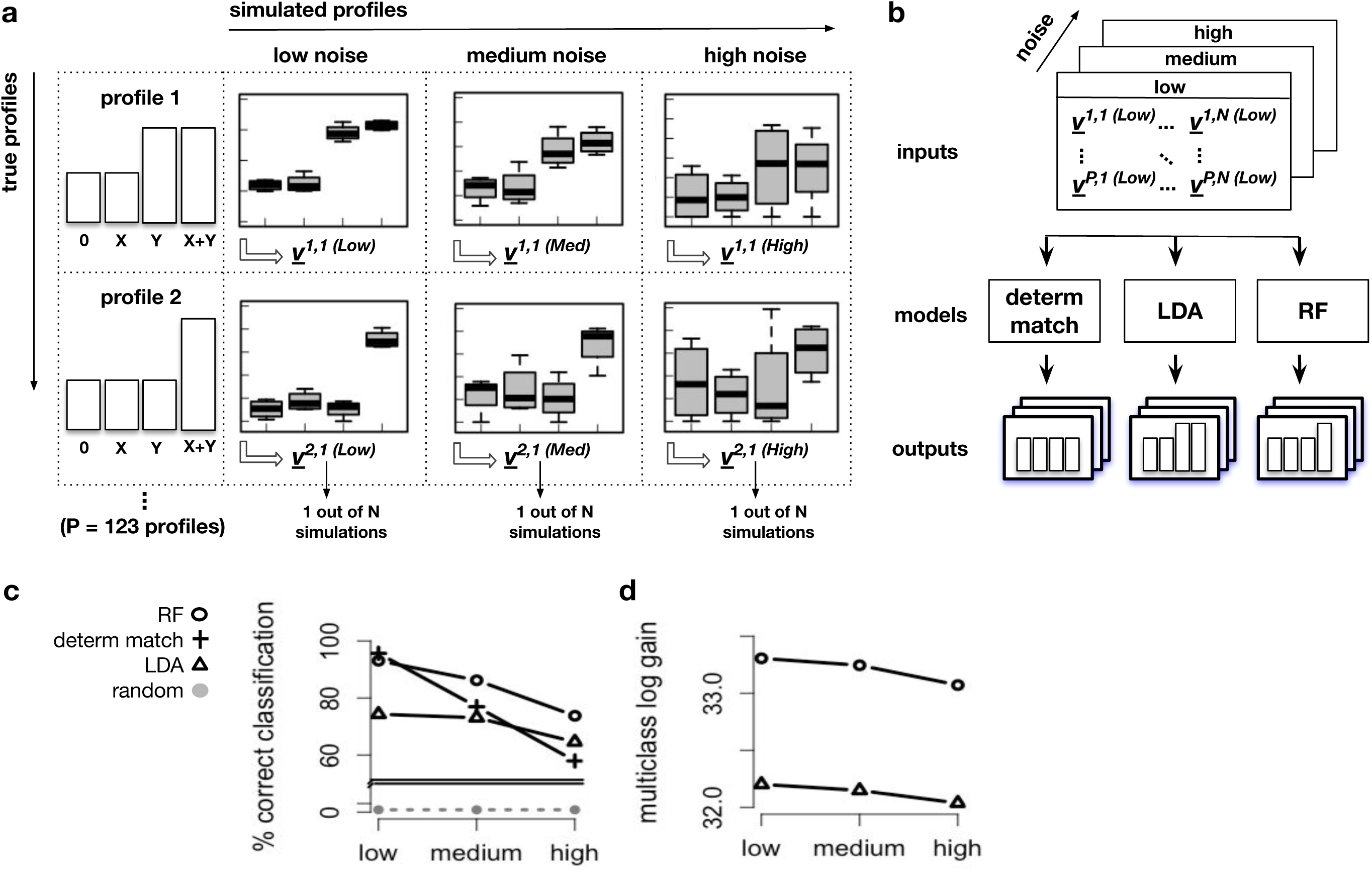
Machine learning classification of treatment interactions. (**a**) We generated a training set of simulated interaction profiles. Each profile was simulated multiple times with realistic group means for the conditions 0, X, Y, X+Y. To build a robust classifier, we simulated three noise levels: low, medium, high. From each simulated profile *i*, instance *j*, and noise level *n*, we extracted a vector ***v***^***i***,***j (n)***^of statistical features including the p-values for all possible pairwise contrasts from the groups 0, X, Y, X+Y. **(b)** The vectors ***v***^***i***,***j (n)***^, labeled with their originating profiles, provided a training set. This was used to develop a machine learning classifier that takes as input a vector of statistical features, and predicts as output the most probable profile. We compared three classification algorithms: Deterministic Match, Linear Discriminant Analysis (LDA), and Random Forests (RF). **(c)** To compare their performance, we measured the classification accuracy -the fraction of correct predictions- on independent test sets. RF provided the most consistent distribution of accuracy. **(d)** To further compare LDA and RF, we computed the multiclass log gain, a metric that accounts for the full probabilistic output returned by these classifiers. Again, RF showed the best performance and was selected as the most robust model.

From each simulated instance of a profile, we extracted a vector of statistical features including the group means, the average deviation from additivity, and the significance of all pairwise contrasts from the conditions 0, X, Y, X+Y (**see Methods**). These features served as predictors of the true class. Overall, the training set comprised ∼340,000 simulated profiles, pairing up vectors of statistical features (inputs) with the corresponding true profile labels (outputs) (**Fig. 2b**). We then trained two established machine learning classifiers: Linear Discriminant Analysis (LDA), and Random Forest (RF), keeping our previously proposed deterministic match algorithm as a reference^6^.

To compare the performance of the different algorithms, we evaluated various metrics for multiclass classification on independent test sets. RF showed a more robust performance then LDA across the three noise levels (**Fig. 2c**). Although the deterministic match had the largest accuracy in the low noise regime, its performance declined more rapidly with increasing noise compared with RF and LDA. An additional advantage of RF and LDA over the deterministic match is the possibility for a “soft” (i.e. probabilistic) assignment of an input profile into any element of the taxonomy. The probabilistic outputs generated by RF systematically showed superior performance over LDA in all noise regimes (**Fig. 2d**). Additional machine learning methods were also evaluated but performed less efficiently in terms of computational overhead.

These results, complemented by analyses of the distribution of precision and recall over all taxonomy classes (**Fig. S2**), indicated RF as the most robust and suitable way to classify combinatorial interactions.

### Building a combinatorial landscape of human immunity

Next, we developed a strategy to systematically map, annotate, and analyze interactions from diverse combination treatments as available in Gene Expression Omnibus (**Fig. 3a**). To retrieve the relevant datasets, we used key terms typically associated with combination treatments such as “synergy”, “antagonism”, “combinatorial”, and similar. This approach was meant to facilitate automatic update of the resource as new *-omics* combination treatments become publicly available. Focusing on immunology, we selected a total of 25 human and 7 murine datasets (**Table 1**). We applied SAIL to each dataset and mapped a total of 29,479 interactions. The proportions of interactions from each type of cell or combination of stimuli varied widely (**Fig. S3**).

**Table 1.**
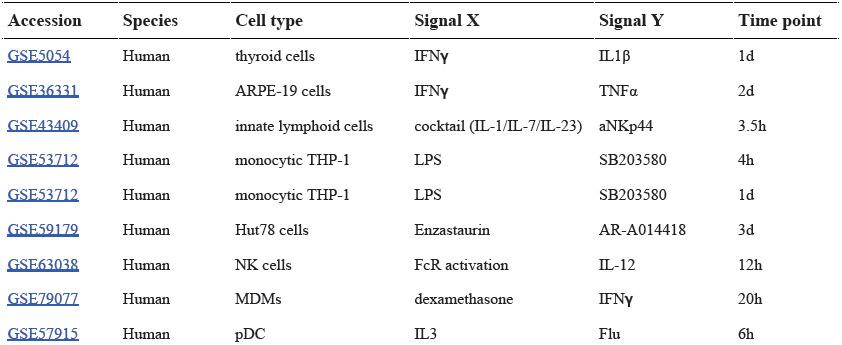
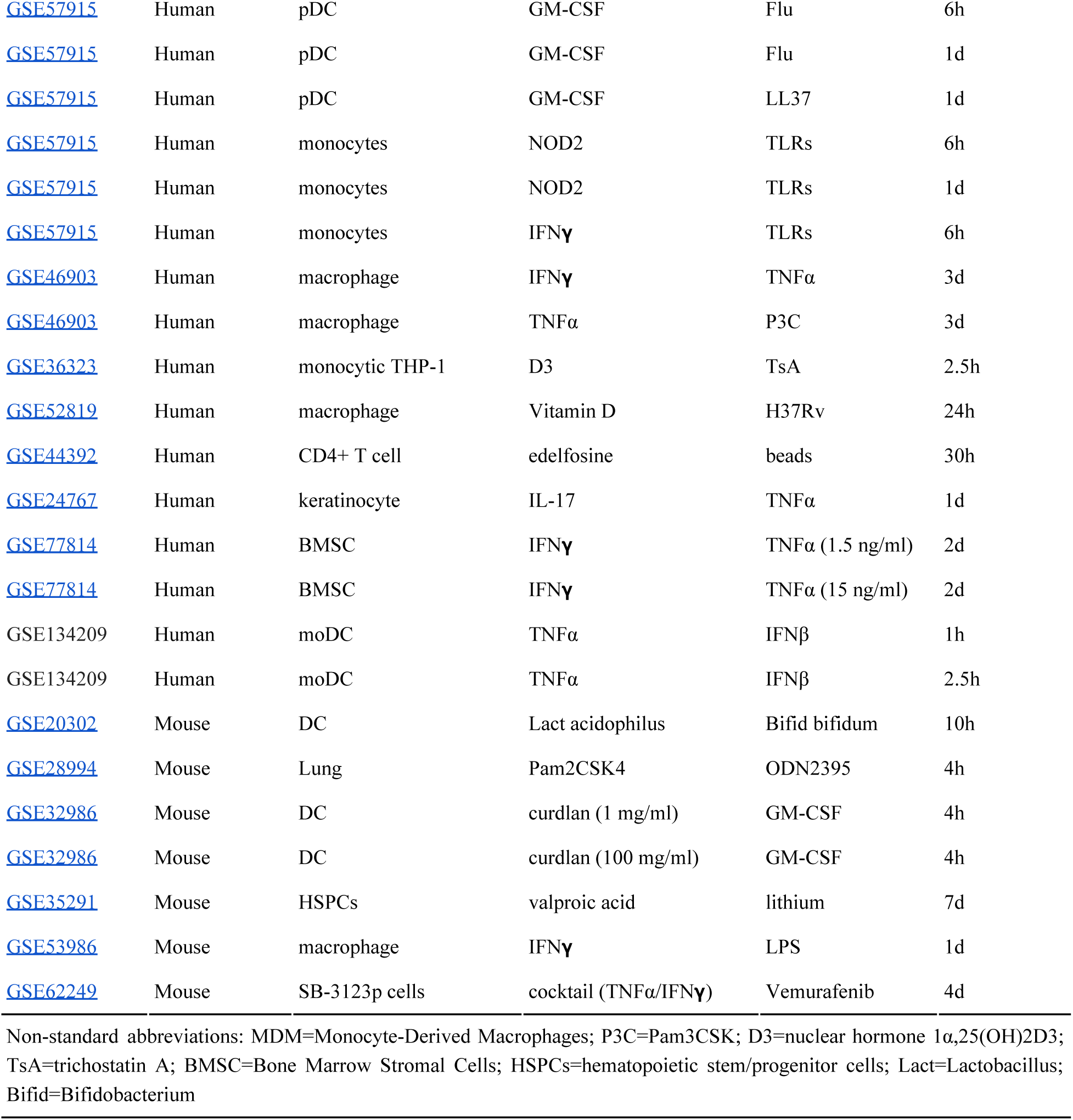
Datasets used to construct the combinatorial landscape of immunity.

**Figure 3.**
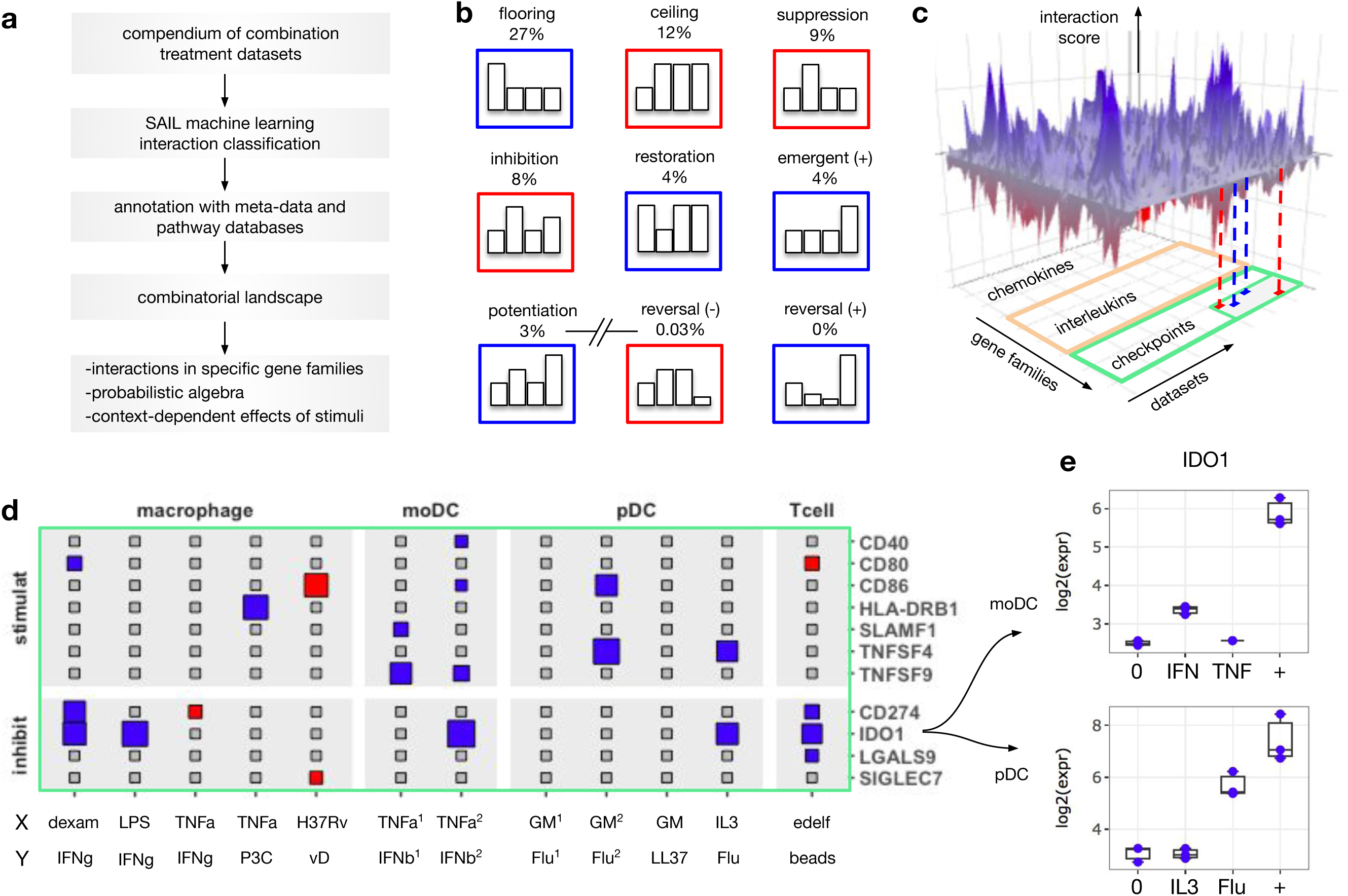
Building a combinatorial landscape of immunity. (**a**) Workflow to map and investigate combinatorial effects from multiple combination treatment experiments. We selected 32 *-omics* combination treatments from diverse immune cells and combinations of stimuli. Applying SAIL to each dataset, we mapped ∼30000 interaction effects. **(b)** The first seven cards represent the most frequent types of interactions. The last two cards represent interactions that occur with vanishingly low frequency**. (c)** To integrate interactions from different datasets, we created a 3D structure with axes representing datasets, genes, and scores quantifying the intensity and robustness of the effects. The resulting landscape, supplemented with metadata and prior knowledge, makes it possible to comprehensively investigate the effect of combination treatments on immune cells. **(d)** The plane shows a 2D projection of the landscape focusing on immune checkpoints, a key immunological gene family. The size and color of the rectangles keeps track respectively of the interaction score and sign (blues: synergistic, red: antagonistic). The immune checkpoint *IDO1* is synergistically induced in multiple datasets (non-standard abbreviations: P3C= Pam3CSK; vD=vitamin D; GM=GM-CSF; edelf=edelfosine; Flu=influenza virus). **(e)** By looking at the specific nature of these synergies, we found two cases of potentiation in human moDC (**top**) and pDC (**bottom**).

Despite numerous studies on combination treatments, the frequency at which different types of interactions occur has not been systematically studied. We found that the most frequent interactions are interpretable as technical and/or biological saturation of the assay, which we refer to as floor and ceiling effects (**Fig. 3b, top subpanel)**. Our approach segregates these effects from less frequent but more biologically relevant interaction responses. The most prevalent of these more important profiles are suppression (9%), inhibition (8%), restoration (4%), emergence (4%), and potentiation (3%) (**Fig. 3b, middle subpanel**). Notably, our analysis also revealed that several theoretically possible combinatorial effects were nearly absent (<0.04%). The rare patterns include reversals, where two signals with the same individual effect (e.g. up-regulation of a gene separately by X and Y) are reversed by the combination (e.g. down-regulation of the same gene by X+Y) (**Fig. 3b, bottom subpanel)**.

To assemble results from multiple combination treatments, we created a 3D landscape with axes representing datasets, genes, and scores quantifying the intensity and robustness of the identified interactions (**Figure 3c**). The dataset axis was also annotated with metadata on the experiments including species, cell type, stimuli, and time point. The gene axis was annotated with immunological gene families such as chemokines, interleukins, checkpoints and other terms from the ImmPort database ^8^. The interaction axis was annotated by the profiles predicted for each interaction by the machine learning classifier.

Slicing the landscape along specific dimensions provides different types of insight into the role and functions of the interactions. For example, slicing by gene family allows systematic identification of synergistic and antagonistic effects involving immune modulators of interest. **Figure 3d** shows a 2D projection of the landscape that contains stimulatory and inhibitory checkpoints, key regulators of the immune system with increasing therapeutic applications ^9,10^. In the considered datasets, immune checkpoints show a variable propensity towards synergistic and antagonistic regulation. While *CD40* and *CD80* present sparse, selective interaction effects across datasets, *IDO1* is synergistically induced in a diversity of datasets. Among these synergies, we found two cases of potentiation in pDC and moDC (**Figure 3e**). Because *IDO1* inhibits T cell division and promotes regulatory T cells ^11^, these synergistic effects may serve to contain overreacting immune responses in the inflammatory microenvironment.

SAIL ability to integrate results from multiple datasets makes it possible to globally estimate the frequency of different synergistic and antagonistic, and to investigate the impact of these effects on gene families and pathways of interest.

### Probabilistic algebra underlying immune cell responses to combination treatments

If the cellular response to a combination was merely additive, the effect of the combination of stimuli would be uniquely determined as the sum of the individual effects. However, interaction effects open the possibility for diverse synergistic and antagonistic scenarios. By integrating results from multiple *-omics* datasets, SAIL enables to explore whether any generalized logic rules link the individual to the combined effects of two signals.

To address this problem, we implemented a new analytical approach and summary visualization (**Fig. 4a**). First, we aggregated profiles in our taxonomy that share the same pattern of individual effects, regardless of their combined effect. For example, we grouped interaction profiles for which neither X nor Y have any effect in isolation (**Fig. 4a, top panel**). Next, we considered all the possible combinatorial effects. Given that X and Y have no isolated effect, the combination X+Y can produce three qualitative outcomes: up-regulation (synergistic), no effect (additive), or down-regulation (antagonistic). Suppose the three possibilities occur with a frequency of 30%, 50%, 20% among the genes classified in this aggregated profile group, respectively (**Fig. 4a, middle panel**). To represent these frequencies in a compact yet informative manner, we used horizontal bars in the column corresponding to the condition X+Y. The color of the horizontal bars keeps track of the sign of the interaction (blue: synergistic, red: antagonistic, gray: additive) (**Fig. 4a, bottom panel**). We refer to this visualization scheme as a “strata plot”. For each strata plot, we quantified the uncertainty in predicting the combinatorial effects for the given individual effects using a normalized Shannon entropy (**see Methods**).

**Figure 4.**
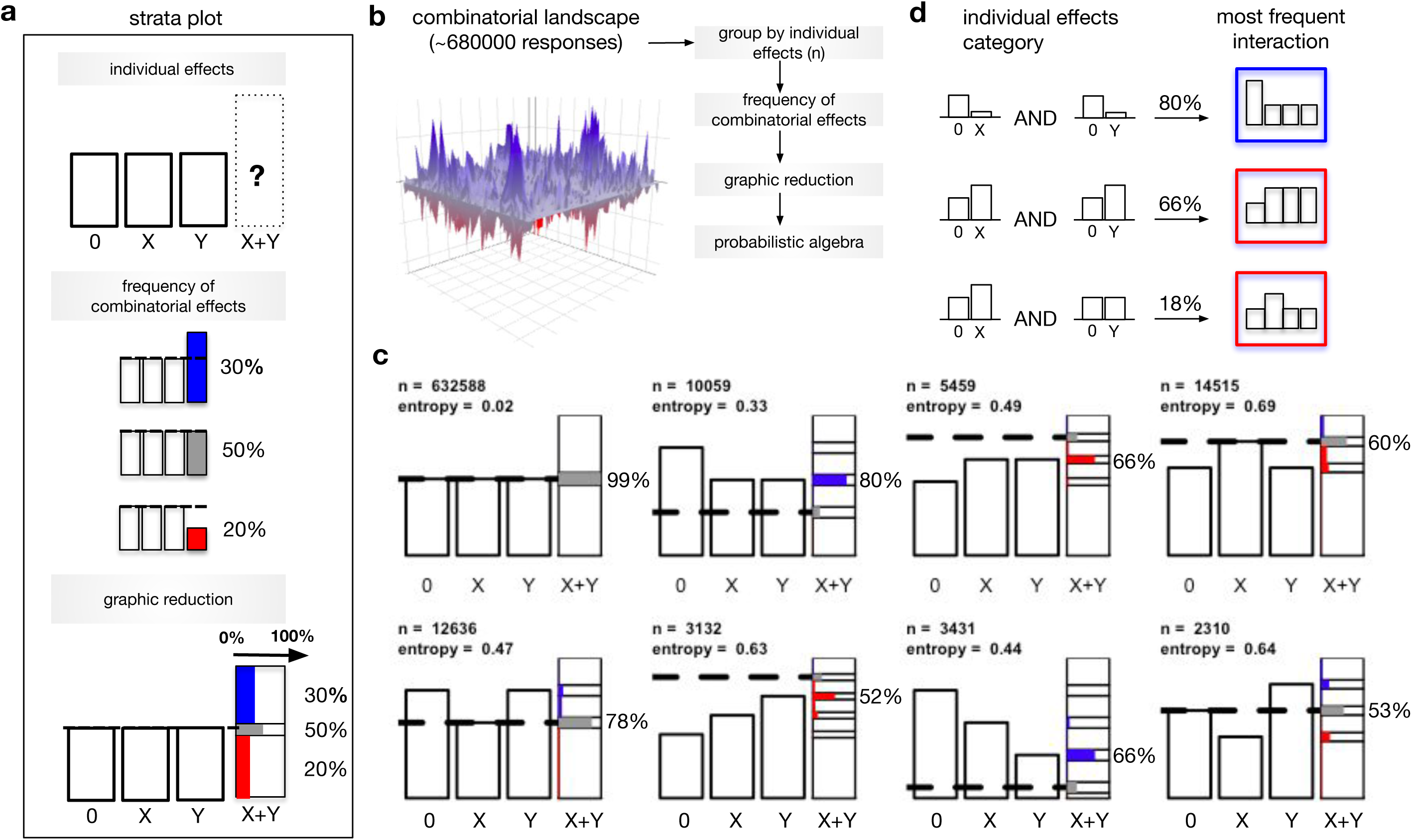
Probabilistic algebra governing immune cell responses to combination treatments. **(a)** To explore the relationship between the individual and combined effects of two signals, we developed a new analytical and visualization technique. First, we divided the taxonomy of response profiles in groups whose elements share the same conjunction of individual effects by X and Y, regardless of their combined effect. For example, Group 1 contains response profiles for which both X and Y have no effect in isolation (**a, top**). Next, we consider all the possible effects of the combination). In Group 1, the combination can have three qualitative scenarios **(a, middle).** We then estimate the frequency of each of these scenarios. In this hypothetical example, the three scenarios are observed with frequencies of 30%, 50%, and 20%. To represent the three frequencies in a compact yet informative manner, we use horizontal bars in the column corresponding to the condition X+Y. We refer to this visualization scheme as a strata plot **(a, bottom)**. The colors of the horizontal bars keep track of the sign of the interaction (blue: synergistic, red: antagonistic, gray: additive). **(b)** By repeating the above procedure for all combinatorial responses in the landscape, we derived a probabilistic algebra linking any conjunction of individual effects. **(c)** Frequency of combinatorial effects corresponding to different types of individual effects. (**d**) Three examples of a probabilistic link between a conjunction of individual effects and the most recurrent interaction.

We then systematically assessed the frequency of all possible combinatorial effects as a function of the individual effects (**Fig. 4b**). For each aggregated profile group, we generated the corresponding strata plot (**Fig. 4c**). The analysis revealed a probabilistic algebra that associates the individual effects of the two signals with the most prevalent type of interaction. For example, if both X and Y downregulate the expression of a gene, the most likely type of interaction is a floor effect (**Fig. 4d, top**). We observed this pattern in 80% out of over 10000 responses in the corresponding group. Similarly, up-regulation individually by both X and Y often results in a ceiling effect pattern in the combination treatment condition, with a frequency of 66% (**Fig. 4d, middle)**. If one signal upregulates a gene, and the other has no effect, the most likely resulting response is an additive effect, followed by a suppression, observed with a frequency of 18% (**Fig. 4d, bottom**). Importantly, this result showed that given two signals, with one up-regulating a given gene and the other having no effect on the same gene, antagonistic effects such as suppression were more frequent than a potentiation effect.

Together with the most likely outcome for each group, our results also reveal the systematic absence of theoretically possible combinatorial effects. Consistent with our analysis of the frequency of the different interactions, we found vanishingly small probabilities associated with reversals.

Altogether, our results support the derivation of a probabilistic algebra underlying the cellular response to combination treatments. The algebra predicts the most likely type of interactions given the individual effects of two stimuli.

### Interactions determine a context-dependent TNFα biology

A critical consequence of interactions is the potential modification of effects observed with each individual treatment. Our framework systematically detects qualitative changes in the effects of a given signal in the presence of other stimuli. To illustrate this, we focused on TNFα, an extensively studied immunomodulators ^12^.

To explore how the composition of the inflammatory microenvironment can alter TNFα biology, we sliced the combinatorial landscape along two axes (**Fig. 5, top-left**). From the dataset axis, we selected combination treatments involving TNFα with other stimuli, including IFNβ and IFNγ in four human cell models (**Fig. 5a, left-margin**). From the interaction axis, we extracted interaction profiles that encoded a qualitative change of the TNFα effect when considered as a mono-treatment. We started by considering three types of qualitative changes: suppression, antagonistic reversal, and synergistic reversal of TNFα effects (**Fig. 5a, top-margin**). For each pair of dataset and profile, we processed the corresponding gene list with enrichment analysis to gain insight at the functional level.

**Figure 5.**
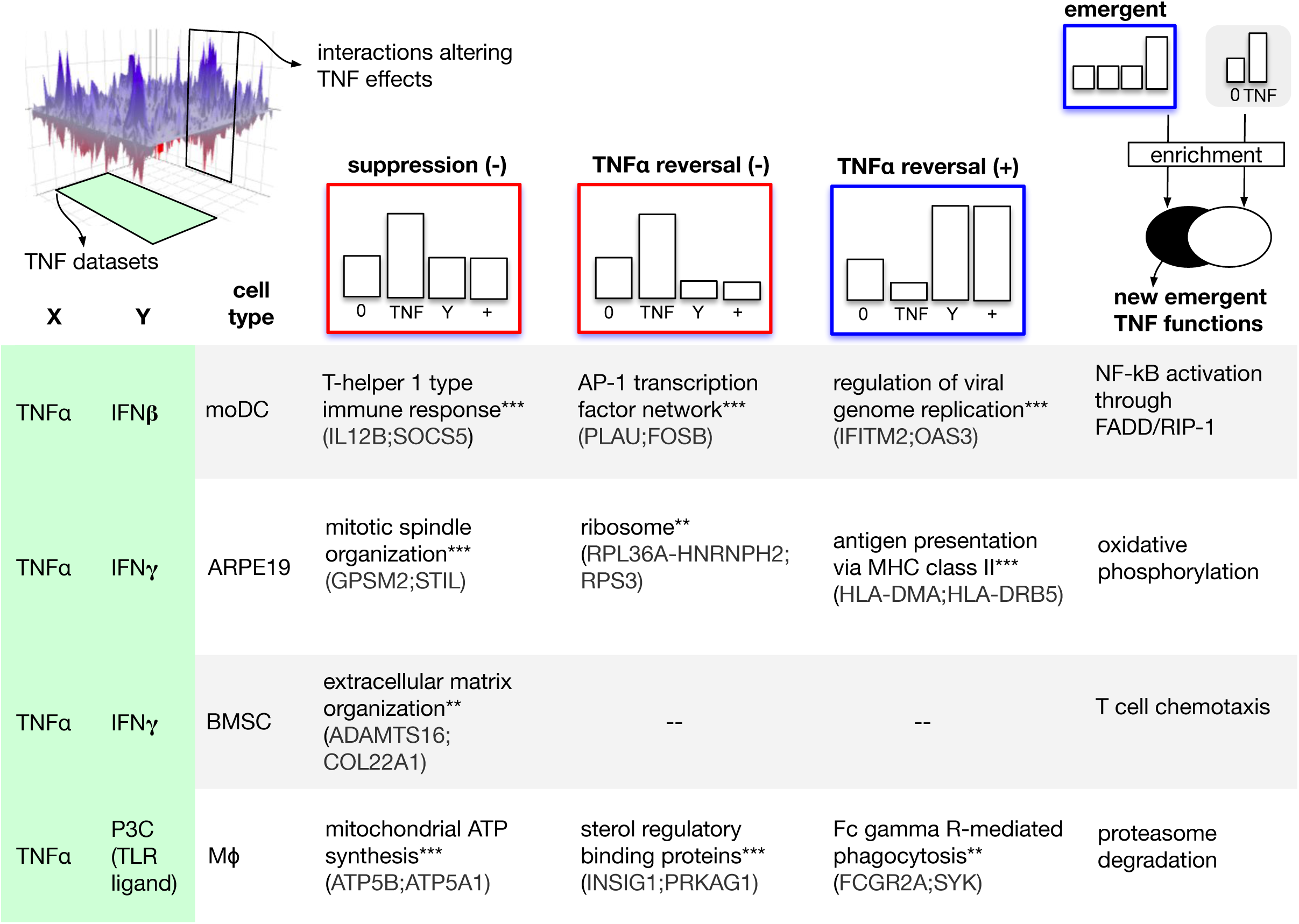
Combinatorial interactions determine a context-dependent TNFα biology. To explore how cofactors might alter the TNFα biology, we sliced the combinatorial landscape (**top-left)** along two axes. From the dataset axis, we extracted combination treatments involving TNFα and concomitant factors including IFNγ and IFNβ in four human cellular models (**left-margin**). From the interaction axis, we extracted interaction profiles that encode a qualitative change of the effect of TNFα mono-treatment. We started by considering three types of qualitative changes: suppression, antagonistic reversal, and synergistic reversal (**top-margin**). For each dataset and profile, we processed the corresponding gene list with enrichment analysis to gain insight at functional level. The matrix elements correspond to selected significantly enriched functions (* adjusted p<0.05, ** adjusted p<0.01, *** adjusted p<0.001). Example hits from each function are shown in parentheses. In the case of emergent effects (last column), we looked for a significant enrichment in new functions, not observed with TNFα alone (**top right**, see also **Methods**). This analysis suggests that cofactors could drive the emergence of new TNFα functions.

Genes showing suppression and reversal of TNFα effects were significantly enriched in important immune processes including T-helper 1 polarization and antigen presentation (**Fig. 5a)**. Further analysis (**Fig. 5, top-right, Methods**) also suggested that co-modulators can drive the emergence of new functions, not observed by TNFα modulation in isolation. Although these emerging functions were relatively few, they comprised potentially important processes such as proteasome degradation and T cell chemotaxis.

Our results illustrate the ability of SAIL to systematically study how the effect of a stimulus is qualitatively altered by other stimuli through a variety of interaction effects including suppression, reversal, and the emergence of entirely new functions.

### Prediction and validation of TNFα and IFNβ interaction effects in human monocyte-derived dendritic cells

Next, we applied SAIL to investigate the synergistic interactions of two specific cytokines, IFNβ and TNFα. While IFNβ and TNFα are key modulators of immune functions whose individual effects have been extensively studied ^12,13^, their interactions remain poorly understood. We previously reported that IFNβ and TNFα act synergistically to induce an antiviral state in monocyte-derived dendritic cells (moDC) ^14^. To investigate the systems level impact of IFNβ and TNFα co-treatment on human moDC, we applied SAIL to the corresponding combination treatment experiment.

SAIL detected 374 synergistic interactions, which we mapped to the corresponding profile groups (**Fig. S4**). The interaction groups with the largest number of synergistic effects were ‘emergent synergy’, ‘TNFα potentiates IFNβ’, ‘IFNβ restores TNFα’, and ‘IFNβ potentiates TNFα’ (**Fig. 6A**). To understand the function of these specific interaction patterns, we sorted the corresponding gene lists based on the synergy score and searched for candidates with potential key immunological roles. We first focused on the emergent synergies due to their special role as responses exclusive to the combination. In this profile, the genes with the largest synergy scores were *LIMK2, MCOLN2, SLC7A5, TP53BP2*, and several genes having more established immunological roles such as *RELB, IL15RA*, and *VCAM1*.

**Figure 6.**
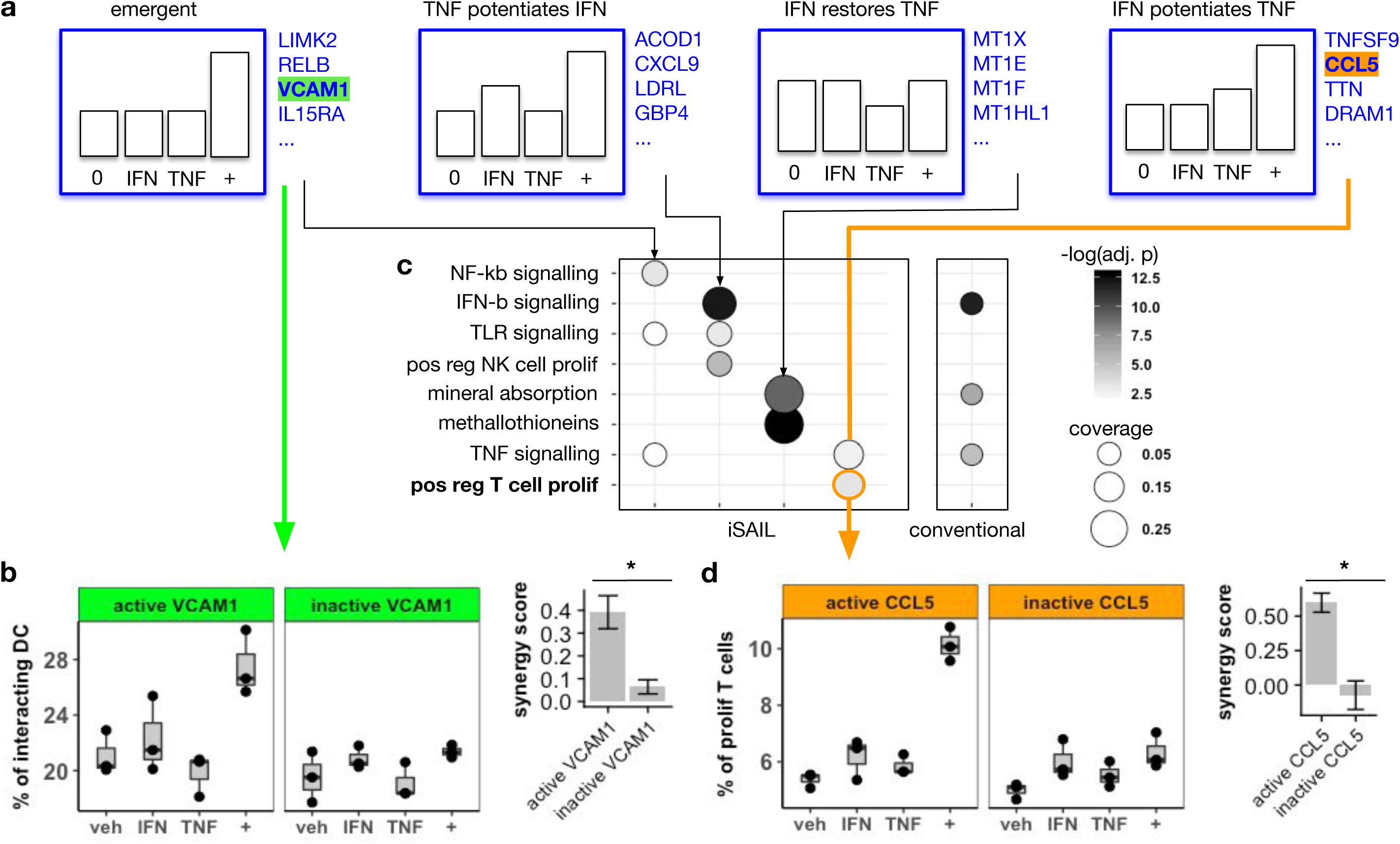
Prediction and validation of IFNβ and TNFα synergistic effects in monocyte-derived dendritic cells. **(a)** We applied SAIL to analyze the synergistic effects induced by IFNβ and TNFα after 1 hour exposure. We focused on the four profile groups with the largest number of synergies. In the group of emergent synergies, we found *VCAM1*, a gene involved in the regulation of cell adhesion. (**b**) We tested the hypothesis that the emergent induction of *VCAM1* in the DC would mediate an increase in DC-T cell interaction. Assayed by imaging flow cytometry, moDCs exposed to IFNβ+TNFα showed an emergent increase in DC-T cell adhesion. This synergistic effect was eliminated by VCAM-1 neutralization. The synergy score for this effect-defined as the mean deviation from additivity- was significantly reduced (t-test, p=0.03). The error bars represent the standard error of the mean synergy score. (**c**) To explore whether IFNβ+TNFα synergy patterns represented coherent gene programs, we determined the functional enrichment for each pattern. (**c, left panel**). We compared these SAIL-based functional analyses to results obtained with a conventional analysis of unclassified synergy genes. (**c, right panel**). SAIL-based enrichment provided a richer functional annotation. In particular, it suggested that synergy genes in the ‘IFNβ potentiates TNFα’ pattern rightmost panel in (**a**) may mediate T cell proliferation. (**d**) This hypothesis was tested using allogeneic cross donor stimulation. The synergy group ‘IFNβ potentiates TNFα’ contained two hits potentially responsible for T cell proliferation: *TNFSF9* and *CCL5*. Using CCL-5 neutralizing antibodies, we confirmed that IFNβ and TNFα act in synergy to promote T cell proliferation, and that this proliferation depends on CCL-5. The synergy score for this effect -defined as the mean deviation from additivity- was significantly reduced by neutralization of CCL-5 (t-test, p=0.008). The error bars represent the standard error of the mean synergy score.

The protein VCAM-1 has been described as a regulator of leukocyte migration and cell adhesion ^15^. Due to the fundamental importance of the DC-T cell axis in the generation of an immune response, we hypothesized that IFNβ and TNFα synergistically induce VCAM-1 to promote moDC-T cell adhesion. We tested this hypothesis by quantifying DC-T cell adhesion using imaging flow cytometry (**see Methods**). When exposed to the combination of IFNβ and TNFα, moDC showed an increased adhesion to T cells that was not observed with either cytokine alone (**Fig. 6d**). The increased DC-T cell adhesion was mediated by VCAM-1, since VCAM-1 neutralization abolished the synergistic effect (**Fig. 6d**). To our knowledge, these results identify for the first time a role for synergistic induction of VCAM-1 by TNFα and IFNβ in promoting DC-T cell adhesion.

To further explore the immune processes controlled by IFNβ and TNFα synergies, we performed enrichment analysis separately for the different profiles, and compared the results with a conventional analysis that aggregates all the synergies in a single gene set (**Fig. 6c, left**). Certain annotation terms were captured by SAIL as well as by the conventional method, but SAIL provided additional insight. For example, both SAIL and conventional analysis captured a highly significant enrichment in mineral absorption. However, SAIL analysis also revealed the pattern ‘IFNβ restores TNFα’ as the main contributor to this enrichment. This pattern contains several members of the family of metallothioneins (*MT1X, MT1E, MT1F, MT1HL1*), which are increasingly recognized as important players in the response to cytokines and pathogen signals ^16^. Overall, annotation by profile revealed an enrichment in annotation terms not resolved by conventional analysis (**Fig. 6c, right**). In particular, we found pattern ‘IFNβ potentiates TNFα’ enriched in T cell proliferation, critical step in the generation of an immune response.

Using an allogeneic cross donor stimulation, we tested the hypothesis that a IFNβ and TNFα stimulation may act in synergy to enhance T cell proliferation (**Fig. 6d, left panel**). The combination treatment induced a nearly two-fold increase in the percent of proliferating T cells, an effect not seen with either stimulus alone. The synergy pattern of the T cell proliferation measurement diverged slightly from the gene level profile, which showed some effect by TNFα alone. It is not surprising that the interactions patterns comparing mRNA level regulation and protein-dependent functional effects show marginal differences. Importantly, the synergistic induction of T cell proliferation, predicted by SAIL, was confirmed experimentally.

Next, we wanted to identify the molecular mediators of the increased T cell proliferation. Candidate genes for mediating this synergistic response predicted by enrichment analysis included *TNSFS9* and *CCL5*. Review of the literature suggested CCL-5 as the most likely candidate ^17^. We therefore hypothesized that CCL-5 may contribute to increased proliferation of T cells induced by INFβ and TNFα exposed DC. This hypothesis was confirmed by immunoneutralization of CCL-5 (**Fig. 6d, right panel**).

These experimental validations demonstrate the value of SAIL in mining interaction data for new hypotheses that guide further study.

## Discussion

In this work, we present a comprehensive machine learning framework to map and interpret interaction effects within and across *-omics* combination treatment studies. Our analysis of a compendium of immunological combination treatment datasets generated a landscape of ∼30,000 interactions. We obtained global insight into the principles and functions of interactions from analysis of the landscape, and validated new hypotheses about combinatorial cytokine effects.

Developing learning models to predict synergistic combinations of treatments based on the individual effects is an active area of research ^18–21^. Despite an apparent methodological similarity, the motivation and goals of our framework are fundamentally different from these studies. In our framework, machine learning is applied to classify and interpret the function of diverse types of synergistic and antagonistic interactions induced by *-omics* combination treatments, and not to directly predict these effects.

Nonetheless, application of SAIL to a compendium of public datasets revealed a combinatorial algebra, that is, a set of rules that for any given pattern of individual effects, can predict the probability of all the possible combinatorial effects. The analysis also revealed that certain *a priori* possible response patterns, such as reversals, are very rare events in all studies examined. This may imply the existence of mechanistic constraints and exclusion principles that limit the spectrum of potential combinatorial responses at the transcriptional level. Overall, our findings could inform and enhance future predictive models with a new type of evidence derived from a number of *-omics* datasets.

A potential consequence of interactions is the radical modification of effects observed with individual treatments. Using the SAIL framework, these events are easily identified by isolating interaction profiles that encode qualitative combination changes in the effect of a treatment of interest. In the case of TNFα, we found that co-modulators alter fundamental immunological processes, such as antigen presentation and T helper cell polarization, and may produce the emergence of entirely new functions, not modulated by TNFα mono-treatment. Identifying the context-dependent effects of an agent may be useful in a therapeutic perspective. The success of therapeutic agents relies on the control of both pathogenic and homeostatic pathways. Our approach may assist in the design of drug combinations that leverage antagonistic interactions to selectively reduce pathogenic activity while preserving necessary homeostatic pathways.

A fundamental tool for biological interpretation of *-omics* experiments is pathway-level enrichment analysis. SAIL uncovers biological processes regulated in combination treatment studies by fine-grained aggregation of genes showing similar interaction responses. As we demonstrate, these aggregates reflect coherent biological processes that provides insight into the principles and functions of interactions, and generate hypotheses for new interaction effects that were experimentally validated. In the analysis of IFNβ and TNFα co-treatment, SAIL uncovered novel synergies that control the DC-T cell interactions and T cell proliferation, both of which are critical immune processes. Importantly, SAIL analysis also suggested specific hypotheses on the molecular mediators controlling these functions, VCAM-1 and CCL-5, which we validated with neutralization experiments.

We note several potential limitations of the SAIL framework. The classifier was trained under the assumption of normally distributed data. While this assumption is commonly held in the analysis of microarray and RNA-seq data upon a suitable transformation, it may not apply to other assays. Under alternative distributional assumptions, our approach would be adaptable to other *-omics* technologies, such as proteomics, metabolomics, and epigenetics data. To demonstrate the usefulness of SAIL, we restricted the initial landscape to immunology. Future releases of the landscape can accommodate additional datasets applicable to all domains of biological research.

To facilitate the application of SAIL, we developed a user-friendly platform. Users can upload experimental data on new combinations, or re-analyze datasets from our curated database. A few simple steps enable the user to identify and interpret the most relevant synergistic and antagonistic interactions. The platform is connected with external resources including ImmPort ^8^, Gene Cards ^22^, and enrichR ^23^, to provide extensive annotation at single-gene and pathway level. SAIL web tools can be used to generate testable hypotheses about the role of combinatorial interactions in driving biological processes.

Notably, treatment interactions are important in clinical therapeutics and side effects. More than 10,000 clinical trials in the United States alone are studying the effects of drug combinations ^1,2,24^. By uncovering relevant interactions and their functions, SAIL can further understanding of interaction mechanisms and the development of combination therapeutics.

## Acknowledgments

This work was supported by the US National Institutes of Health (NIH) grant 5U19AI117873, by the European Research Council under Grant IT-DC 281987, by Agence Nationale de la Recherche under Grant ANR-11-LABX-0043 CIC IGR-Curie 1428, and by ERC 2015 POC DrugSynergy 680890.

## Author contributions

Conceptualization: A.C., E.Z., S.C.S., V.S.; Formal analysis: A.C., E.Z., and S.T.J.; Experiments: B.M.H.; Writing, reviewing, and editing: A.C., E.Z., V.S., and S.C.S.

## Methods

### Definition and simulation of interaction profiles

The notion of interaction profiles has been introduced in our previous work ^6^, and is briefly summarized here. The interaction profiles represent qualitatively different scenarios for the expression of a gene in the conditions 0, X, Y, X+Y. **Fig. S1** shows the taxonomy of 123 profiles used in this study. Mathematically, each profile corresponds to a linear system of equalities and inequalities satisfied by the mean expression levels of a gene in the conditions 0, X, Y, X+Y, respectively denoted by 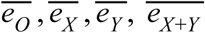. The linear system defining a given profile admits infinitely many solutions, each of which can be seen as a particular instance of the profile. For example, an emergent synergy (no effect by X, no effect by Y, and up-regulation by X+Y) is satisfied by the vectors (2.2, 2.2, 2.2, 5.5), (4.1, 4.1, 4.1, 7.8), as well as by infinitely many other qualitatively similar vectors.

To simulate a profile, our strategy starts by sampling the solution space of the corresponding system of inequalities. The solution space is sampled within a range of admissible values, chosen to mimic the experimental data. Next, a noise term is added to each instance of a profile using a random number generator. The noise term is assumed to be normally distributed. This assumption is widely held in the analysis of microarray data, and still applicable to RNA-seq data upon a suitable transformation ^7^. Increasing noise levels correspond to a decreasing effect size, which is defined in terms of the standardized differences between the group means in the four conditions, as further described below. To account for the relatively small number of samples in *-omics* data, we simulated four replicates for each of the conditions 0, X, Y, X+Y. The steps to simulate interaction profiles are as follows:

#### Definition of a range of admissible expression values

This was chosen as the interval [-14, 14], consistent with the range of log2-transformed expression values from microarray and RNA-seq data upon the Voom transformation.

#### Sampling the solution space for the given profile in the specified range

The vector of numbers 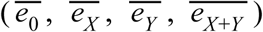 for the given profile were found with the function *xsample* from the package *limsolve*. For each profile, we extracted 400 instances for each level of noise.

#### Definition of the signal of a simulated profile

The signal can be seen as a generalization of the fold-change in A Vs. B experiments. In a combination treatment, we first consider all pairwise fold-changes from the conditions 0, X, Y, X+Y. Except for the case of constant genes, at least one of these contrasts must be different from 0. The signal δ is defined as the absolute value of the smallest non-zero difference. To avoid very weak signals, not meaningful in the analysis of expression data, a minimum signal of 0.5 is used in the training set.

#### Simulation of random noise

To simulate random variability around the values 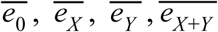, the data was assumed to be normally distributed around the group means: 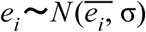, with *i = 0, X, Y, X+Y*. The parameter σ was assumed the same for all groups. For each of the four conditions, we simulated 4 replicates. Different levels of noise, were simulated by setting different values of the ratio δ/σ.

#### Enforcement of the range of the expression values

The addition of random noise can push some of the values *e*_*i*_ outside the initially prescribed range of expression. In this case, we forced the simulated values to be at the limit of the range. For example, a value of −18.5 was reset to −14, the lower limit of the prescribed range.

### Training and testing of the machine learning classifier

To train the machine learning classifiers, we generated a training set by simulating multiple instances of every profile in the admissible range of expression values. For each instance of a simulated profile, we extracted a vector of statistical features which were used as predictors of the true class. The statistical features were built on the output of the *Limma* package, an established tool for differential gene expression analysis ^25^. These include the estimated mean Values 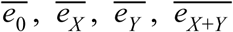, the p-values of all possible pairwise contrasts among these four values, and additional statistics returned by *Limma*. The training set comprised different noise regimes. These were simulated by fixing different values for the parameter δ/σ, as described in the previous section. We considered the following values: low noise (δ/σ =4), medium noise (δ/σ =2.5), high noise (δ/σ =2). Training of the machine learning classifiers was done using the R packages *Caret* and *RandomForest*.

To select the best model, we generated additional simulated data and measured the out-of-sample classification accuracy per profile and for different values of δ/σ defined as in the training set. For each of these values, the accuracy was quantified as the proportion of correct predictions. The multiclass log-gain, and the class-specific precision and recall were computed with the packages *MLmetrics* and *mltest.*

### Generating the combinatorial landscape of immunity

Public combination treatment datasets were retrieved from Gene Expression Omnibus using the package *GEOquery* ^*26*^. To facilitate comparisons, all the datasets were imported in the same format as in the original publications. The datasets were preprocessed as follows. First, if different probes were available for the same gene, the probe with the largest coefficient of variation was selected. Second, genes with low coefficient of variation (lower than the median of the distribution computed for all genes) were filtered out. Next, differentially expressed genes were determined with the *Limma* package. A significance cutoff of 0.05 was applied on the p-values after correction for multiple testing. An additional cutoff was imposed on the δ (defined above): genes with δ lower than the median value computed across all differentially expressed genes were filtered out. The resulting differentially expressed genes were then analyzed by the machine learning classifier which assigned to each gene the predicted element of the taxonomy. For each identified interaction, a score was defined to measure its magnitude as well as its significance. The magnitude of the interaction *b* was measured as the average Bliss index, defined as the average deviation from additivity 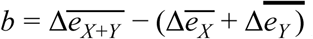, where 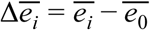, with *b* > 0 for synergistic effects and *b* < 0 for antagonistic effects. The significance of the interaction was measured as the class probability *p* returned by the classifier. The overall score was defined as the product *b* · *p*. The identified interactions were annotated using a manually curated list of stimulatory and inhibitory immune checkpoints, as well as the gene lists provided by the ImmPort database ^8^.

### Probabilistic algebra of signal integration

To derive the probabilistic algebra, we first organized the taxonomy of profiles in groups. Each group consists of profiles that share the same individual effects of X and Y, but differ for the effect of the combination X+Y. Because the labels X and Y are arbitrary and biologically irrelevant, pairs of groups that obtained by switching the role of X and Y were considered as the same group, for a total of 10 groups.

Let *N*_*i*_ denote the number of responses classified in group *i* from all analyzed datasets, for *i* = 1, …, 10. For each of these groups, let *n*_*i*_ denote the number of possible outcomes for the combination. Furthermore, let *N*_*i, j*_ the number of responses in group *i* with outcome *j*, for *j* = 1, …, *n*_*i*_. The probability of outcome *j* in group *i* is estimated as *p*_*i,j*_ = *N*_*i,j*_/*N*_*i*_. Given the probabilities *p*_*i,j*_, the normalized Shannon entropy for group *i* was computed as 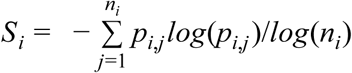. This is a number between 0 and 1 that measures the uncertainty in the response to the combination for given individual effects. A value of *S*_*i*_ = 0 corresponds to a deterministic response (only one outcome for the combination observed with 100% frequency), while *S*_*i*_ = 1 corresponds to maximum uncertainty (all outcomes for the combination occur with 1/*n*_*i*_ frequency).

### Enrichment analysis

The functional enrichment of interactions was done using the Enrichr library ^23^. Four annotation databases were considered: GO Biological Processes (2017b), KEGG (2016), Wikipathways (2016), Reactome (2016). Enrichment was considered significant if the enrichment p-value adjusted for multiple testing was lower than 0.05.

To analyze the synergies induced by IFNβ and TNFα co-treatment, we focused on annotation terms with size lower than 500 genes, to increase the specificity of the identified functions and pathways. Moreover, we imposed a minimum threshold in the overlap between the annotation term and the gene set being analyzed. This threshold was meant to identify annotation terms covering a minimum proportion of the gene set being analyzed. We chose a minimum coverage of 2%.

### DC differentiation

All human subjects research protocols were reviewed and approved by the IRB of the Icahn School of Medicine at Mount Sinai. Monocyte-derived DCs were obtained from healthy human blood donors following a standard protocol described elsewhere ^27^. All experiments were replicated using cells obtained from different donors. IFNβ and TNFα treatments IFNβ (PBL InterferonSource) was added at a concentration of 2000 U/ml and TNFα (Symansis) at a concentration of 1.3 ng/ml to the DC culture. Incubation time varied.

### Microarray data of human moDC treated with IFNβ and TNFα

DC were treated with 4500 pg/mL TNFα, 3000 pg/mL IFNβ, or the combination of both for either 1 h or 2.5 h. Untreated DC served as a negative control. Three samples were taken per treatment and time point. RNA was extracted with the RNeasy plus kit (Qiagen) following the manufacturer’s instructions. Gene expression was assayed using broad human genome specific HG-U133_Plus_2 GeneChip expression probe arrays (Affymetrix). Affymetrix microarray data were normalized using gcRMA ^28^. Additional data processing was done as steps described above (see section “Generating the combinatorial landscape of immunity”).

### Experimental validation of IFNβ and TNFα synergistic effects

To test the involvement of VCAM1 on TNFα IFNβ induced synergy, DCs were exposed to TNFα, IFNβ, the combination of TNFα and IFNβ or control as described above. Four hours after treatment DCs were exposed to allogeneic T cells in a 1:3 ratio for an additional four hours and then fixed with paraformaldehyde. Cells were stained with fluorochrome labeled antibodies against CD11c (DCs) and CD3 (T cells) and analyzed by imaging flow cytometry. DCs interacting with T cells were identified in images were one or multiple T cells had a direct contact with a DC.

To test the involvement of CCL5 on TNFα and IFNβ induced synergy, DCs were exposed to the cytokine mixtures as described above. After 4 hours, DCs were exposed to CFSE stained allogeneic T cells for 5 days and then fixed with paraformaldehyde. Cells were stained with a monoclonal antibody against CD11c and the extend of T cell proliferation was measured by the dilution of CFSE in the CD11c negative population, as CFSE gets weaker with every T cell division. The stain to exclude DCs was necessary as DCs also digest CFSE positive parts of T cells.

## Data and code availability

All the analyses have been implemented in R. An interactive R Shiny application of SAIL can be found at https://SAIL.shinyapps.io/test_app/. The site also contains downloadable code and documentation to run the software locally.

## Supplemental information

**Figure S1.**
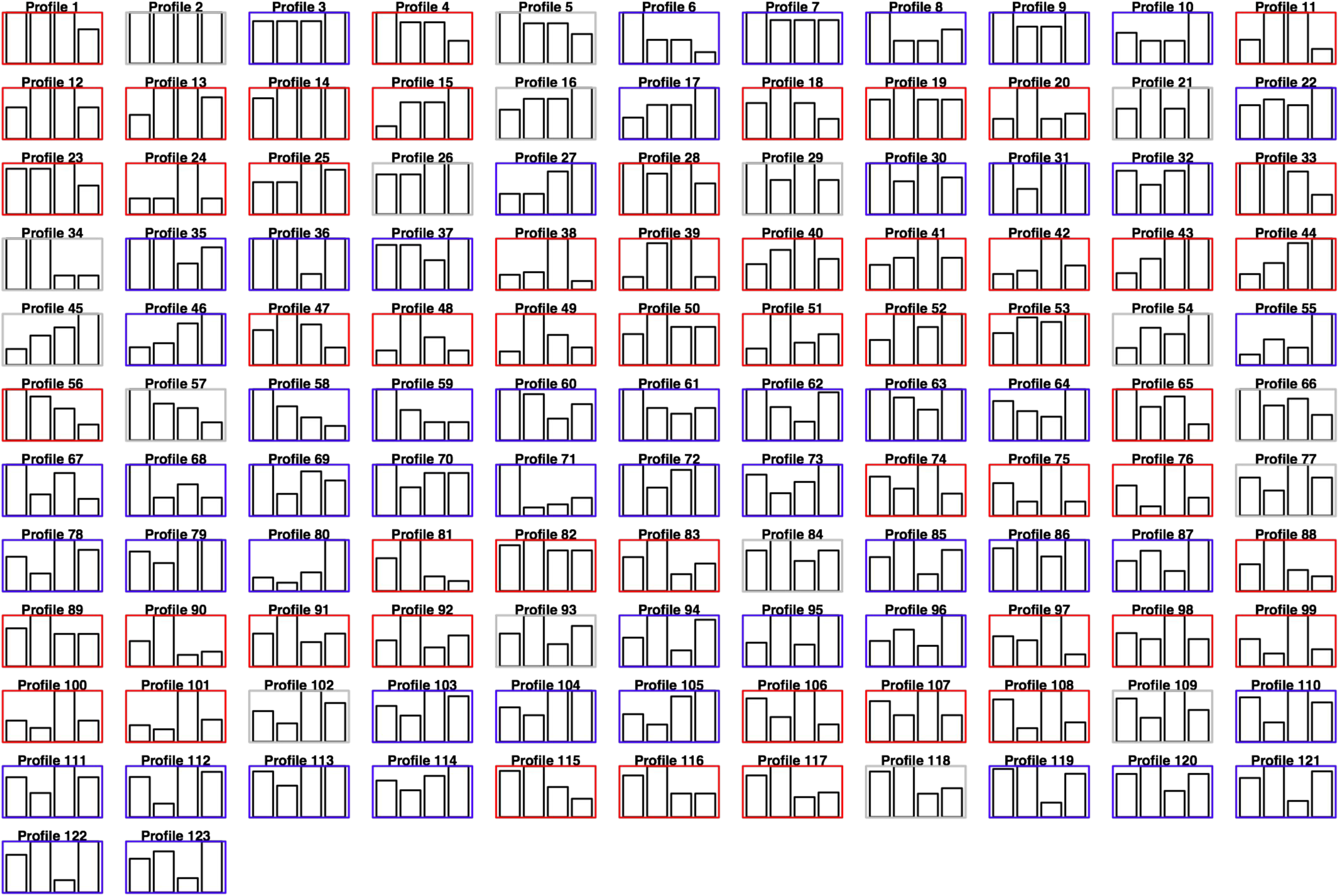
A comprehensive taxonomy of response profiles to combination treatments. Each card corresponds to a possible response profile of a standard combination treatment experiment involving the conditions 0 (control), X, Y, and the combination X+Y. The color code keeps track of the sign of interaction: gray for additive, red for antagonistic, blue for synergistic. The taxonomy includes all the qualitative responses defined in our previous work^6^, as well as additional profiles that capture the semi-quantitative effect of X, Y in case these two signals have opposite effects (e.g., one gene is up-regulated by X and down-regulated by Y or viceversa). In this case, we introduce new profiles by comparing the magnitude of the opposing effect.

**Figure S2.**
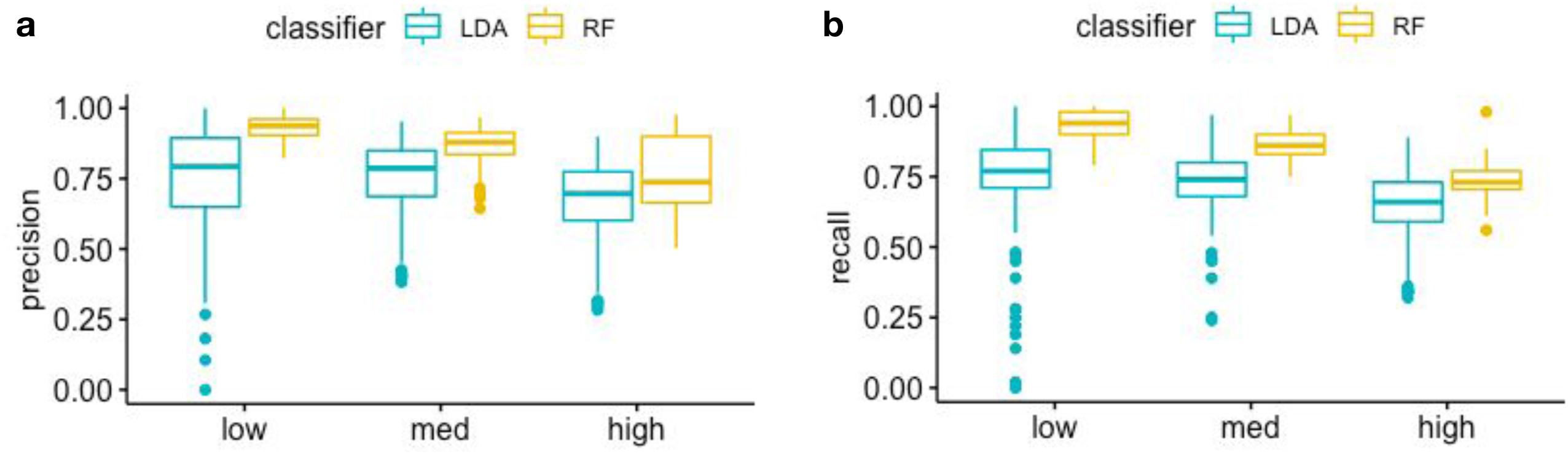
Distribution of precision and recall of LDA and RF across the taxonomy of response profiles. To evaluate the performance of LDA and RF, we measured the distribution of precision (**a**) and recall (**b**) over all the 123 elements in the taxonomy of response profiles to combination treatments (see **Figure S1**). These metrics were evaluated on independent test sets and for three levels of noise: low, medium, and high. The results show that LDA fails to detect certain response profiles even in the low noise range. Overall, RF provides higher precision and recall, and a more consistent performance over the different classes.

**Figure S3.**
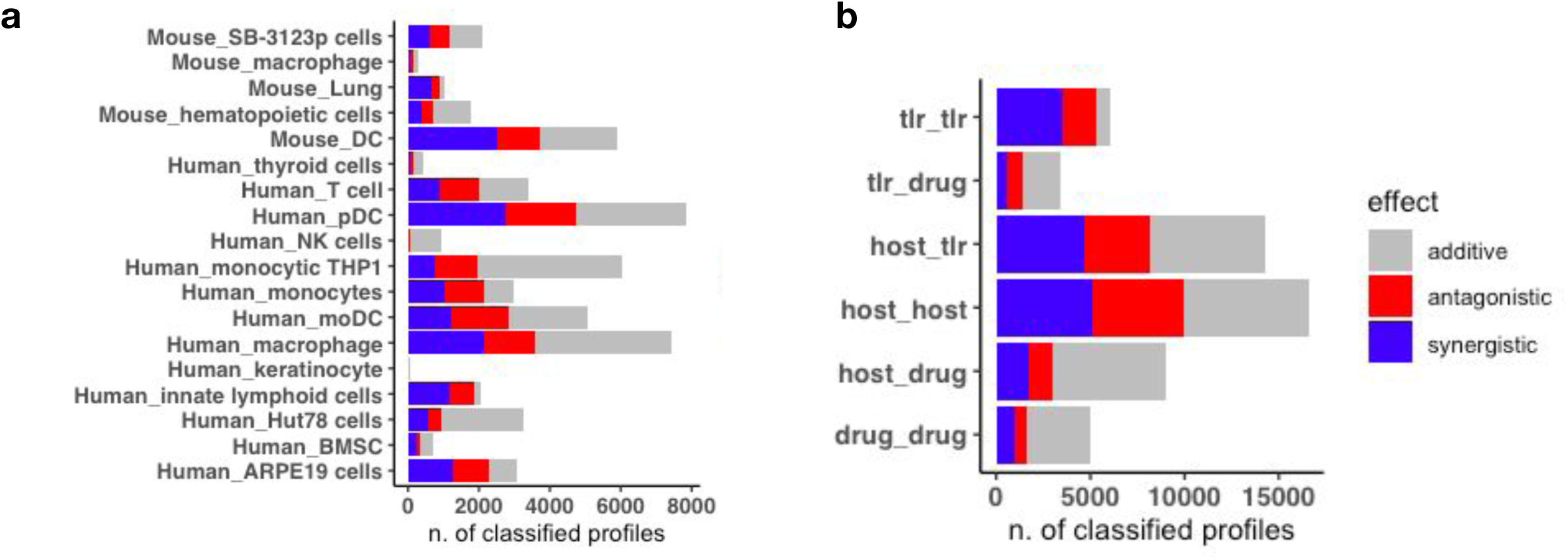
Quantification of interaction by cell type and type of combination. Quantification of the ∼30000 interactions included in combinatorial landscape aggregated by immune cell type (**a**) and by type of combination (**b**). Non-standard abbreviations: tlr=TLR ligands; host=host-derived factors (e.g. cytokines).

**Figure S4.**
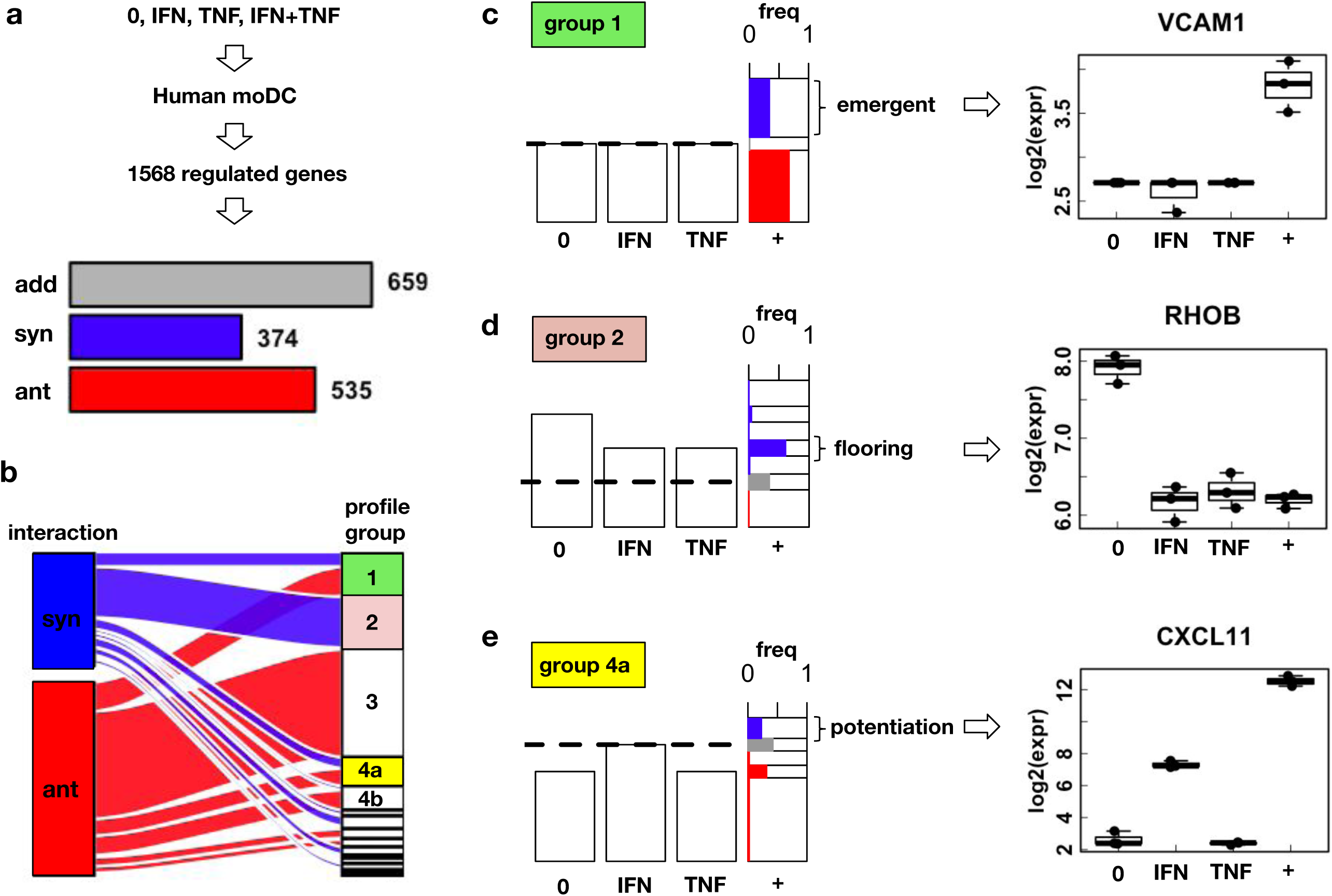
SAIL analysis of interaction effects between IFNβ and TNFα in human moDC. (**a**) To study the interactions of IFNβ and TNFα, human blood moDC were treated in triplicates with control (0), IFNβ, TNFα, and the combination IFNβ+TNFα. After one hour, gene expression was measured with Microarray chips. Differential expression analysis revealed 1568 genes (adjusted p-value<0.05). (**a, bottom**) Using SAIL, these genes were classified in 596 additive, 359 synergistic, and 613 antagonistic profiles, and (**b**) mapped the corresponding profile groups. (**c**) Profile group 1 contains emergent effects, i.e., genes regulated only with the combination, as exemplified by the gene *VCAM1* (**c, right sub-panel**). (**d**) Profile group 2 contains synergies interpreted as a floor effect, as exemplified by the gene *RHOB* (**d, right sub-panel**). (**e**) Profile group 4b contains genes up-regulated by IFNβ and further potentiated by the combination. This effect, exemplified by the gene *CXCL11* (**e, right sub-panel**), is interpreted as a ‘potentiation’ effect.

